# Sidestream bio-P and mainstream anammox in a BNR process with upstream carbon capture

**DOI:** 10.1101/2023.04.11.536442

**Authors:** Kester McCullough, Stephanie Klaus, Christopher Wilson, Peter A. Vanrolleghem, April Z. Gu, Charles Bott

**Affiliations:** modelEAU, Département de génie civil et de génie des eaux, Université Laval, 1065 av. de la Médecine, Québec, QC G1V 0A6, Canada; Hampton Roads Sanitation District, 1434 Air Rail Avenue, Virginia Beach, VA, 23455, USA; School of Civil and Environmental Engineering, Cornell University, 220 Hollister Hall, Ithaca, NY, 14853, United States

**Keywords:** biological phosphorus removal, A/B process, carbon capture, shortcut nitrogen removal, anammox

## Abstract

The integration of biological phosphorus removal (bio-P) and shortcut nitrogen removal (SNR) processes is challenging because of the conflicting demands on influent carbon: SNR allows for upstream carbon diversion but this reduction of influent carbon (especially volatile fatty acids (VFA) prevents or limits bio-P The objective of this study was to achieve SNR, either via partial nitritation/anammox (PNA) or partial denitrification/anammox (PdNA), simultaneously with biological phosphorus removal in a process with upstream carbon capture. This study took place in a pilot scale A/B process with a sidestream bio-P reactor and tertiary anammox polishing. Despite low influent rbCOD concentrations from the A-stage effluent, bio-P occurred in the B-stage thanks to the addition of A-stage WAS fermentate to the sidestream reactor. Nitrite accumulation occurred in the B-stage via partial denitrification and partial nitritation (NOB out-selection), depending on operational conditions, and was removed along with ammonia by the tertiary anammox MBBR, with the ability to achieve effluent TIN less than 2 mg/L.

## 1 Introduction

Partial nitritation/anammox (PNA) in sidestream processes, treating dewatered anaerobically digested sludge liquor, is well established with over 100 full-scale installations (Lackner et al., 2014). The main challenges of achieving mainstream deammonification are NOB out-selection and anammox retention (Cao et al., 2017). NOB repression is easier in sidestream processes thanks to high free ammonia (FA) concentrations (Anthonisen et al., 1976) and high temperature (Hellinga et al., 1998). The challenge of anammox retention can be overcome by a two-phase, separate SRT, system in which nitrifiers and ordinary heterotrophs are in a suspended growth reactor, followed by a completely anoxic anammox moving bed biofilm reactor (MBBR) (Regmi et al, 2015; Ma et al., 2011). However, sustaining NOB out-selection has proven very difficult, and may be an insurmountable obstacle to full-scale adoption of mainstream PNA (Wells et al., 2017; Ma et al., 2016; Cao et al., 2017; Lotti et al., 2015)

Due to the difficulty of generating nitrite from partial nitritation, recent research efforts have explored the generation of nitrite from partial denitrification (Du et al., 2017; Le et al., 2019; X. Wang et al., 2019). The operational requirements for partial denitrification are far less stringent than those for NOB out-selection, leading to improved feasibility and stability. While partial-denitrification anammox (PdNA) offers less aeration and external carbon savings than PNA, it still provides significant savings over full nitrification/denitrification. PNA theoretically provides at most 60% aeration savings and 100% external carbon saving, while PdNA provides at most 50% aeration savings and 80% external carbon savings, but the realized aeration savings will be dependent on how efficiently influent carbon is used for denitrification and how much influent carbon is available (Ma et al., 2016; McCullough et al., 2022). These processes can further increase aeration savings and reduce energy demands by allowing carbon diversion upstream of the BNR process (Daigger, 2014; McCullough et al., 2022), for example by using a high-rate activated sludge (HRAS) A-stage process (Bӧhnke and Diering, 1997; Miller et al., 2017).

While PdNA has been considered for sidestream treatment (Kalyuzhnyi et al., 2008; Sharp et al., 2017), it has greater potential for mainstream treatment because of the possibility to take advantage of influent carbon for partial denitrification, and PdNA obviates the need for NOB out-selection in the mainstream. The factors that have been reported to impact the occurrence of partial denitrifying microorganisms include the type of carbon sources, limited ratio of carbon to nitrogen (C/N), high pH, high salinity, and the alternating feast-famine regime (Le et al., 2019; Zhang et al., 2020). In sidestream treatment, partial denitrification requires external carbon, and NOB out-selection is not a challenge, so PNA remains the more obvious choice.

Previous literature suggest that the nitrite accumulation could be enhanced by either sub-sufficient carbon supply (i.e. low C/N ratio) or lower electron donor flux rates by internal polyhydroxyalkanoates (PHA) hydrolysis to favor the PDN process (Zeng et al., 2003). Particularly, the identities and roles of specific organism that are capable of utilizing internal carbon storage to facilitate the PDN process have hardly been investigated. Partial denitrification could be achieved using internally stored carbon, such as PHAs and glycogen, in which case no external carbon would be required (if influent VFA is the source of the stored compounds). There are some examples of volatile fatty acids in the influent being stored in an anaerobic zone, and then used for denitrification in a post-aeration anoxic zone (Winkler et al., 2011; Vocks et el., 2005; Alleman and Irvine, 1980), or during intermittent aeration (Zhao et al., 1999). This can be carried out by denitrifying PAO (dPAO), denitrifying GAO (dGAO), or other heterotrophs (Tsuneda et al., 2006; Rubio-Rincón et al., 2017; van Loosdrecht et al., 1997). It is not clear from the literature what factors would cause stored carbon-derived partial denitrification to nitrite over full denitrification. Recent studies have shown that dGAO may preferentially reduce nitrate to nitrite, and then dPAO reduce nitrite to dinitrogen gas (X. Wang et al., 2019; Rubio-Rincón et al., 2017). While accumulation of nitrite via partial denitrification by dGAO is desired for treatment by anammox bacteria, GAO compete with PAO for VFA, and can lead to deterioration of bio-P performance (Erdal et al., 2003; Lopez-Vazquez et al., 2009). If PdN cannot be achieved using stored carbon, supplemental carbon sources can be used such as acetate, glycerol, and methanol to drive PdN in an anammox MBBR (Klaus et al., 2023).

Sidestream bio-P (also referred to as sidestream enhanced biological phosphorus removal, S2EBPR) processes are configurations in which the influent is bypassed around the anaerobic zone (Barnard et al., 2017). Sidestream bio-P configurations can include RAS fermentation, in which a portion of the RAS (5-10%) is held in an anaerobic reactor with a retention time of one to two days (Barnard et al., 2017; Gu et al., 2019; Onnis-Hayden et al., 2020; Tooker et al., 2016; D. Wang et al., 2019; Vollertsen et al., 2006). Alternatively, the retention time of the sidestream reactor can be considerably shorter (1-4 hours) with the addition of supplemental carbon (since fermentation is not occurring in the reactor), such as primary sludge fermentate (Cavanaugh et al., 2012; Gu et al., 2020; Tooker et al., 2016).

RAS fermentation was initially investigated as a means to decrease sludge production and nutrient loading to the nutrient removal step, compared to primary sludge fermentation (Andreasen et al., 1997; Novak et al., 2007). It has also been shown that sidestream processes result in more stable bio-P performance (Lanham et al., 2013; Onnis-Hayden et al., 2020). Sidestream processes protect the anaerobic zone from wet weather flows, provide VFA from hydrolysis and fermentation, and offer possible suppression of GAO (Vollertsen et al., 2006; Barnard et al., 2017; Li et al., 2020). The enrichment of PAO over GAO may occur because PAO decay more slowly than GAO under long-term anaerobic conditions (Varga et al., 2018; Li et al., 2018).

The goal of this study was to integrate carbon capture and mainstream anammox with sidestream bio-P. Bio-P is achieved by fermenting a portion of the A-Stage WAS in order to generate enough VFA for addition to the sidestream reactor to drive bio-P in the B-Stage. The challenge behind integrating bio-P and SNR in general, is the conflicting use of influent carbon. The goal of shortcut nitrogen removal is to reduce carbon demand for denitrification so that carbon can be diverted. However, if a HRAS A-stage process is utilized for carbon diversion, then there is little to no rbCOD available in the B-stage process for biological phosphorus removal to occur, thus requiring an alternative configuration such as sidestream bio-P. Bio-P cannot occur in the A-stage because the SRT is too short (less than 1 day) for PAOs to grow.

## 2 Methods

### 2.1 Pilot Configuration

This study took place at the A/B pilot located at the Hampton Roads Sanitation District (HRSD) Chesapeake-Elizabeth Plant located in Virginia Beach, VA, USA (Figure 1). The influent was temperature-controlled to 20°C and fed at constant flow rate to the A-stage high rate activated sludge (HRAS) process (45-minute HRT). The A-stage process configuration and process control is detailed in Kinyua et al. (2017). The effluent from the A-stage clarifier fed the B-stage nitrogen removal step (5-hour HRT). The B-stage consisted of 5 reactors in series. The anaerobic selector was 50 Liters (20 minute HRT), was continuously mixed, and covered with ping pong balls to minimize oxygen transfer. The other 4 intermittently aerated CSTRs were 150 Liters each for a total volume of 650 Liters. Following the secondary clarifier, the B-stage effluent fed an anoxic anammox MBBR with an HRT of 2 hours, total volume of 340 Liters, and a 60% fill of K3 media (Anoxkaldnes, Lund, Sweden). The MBBR at times was fed glycerol to induce partial denitrification combined with anammox.

**Figure 1.**
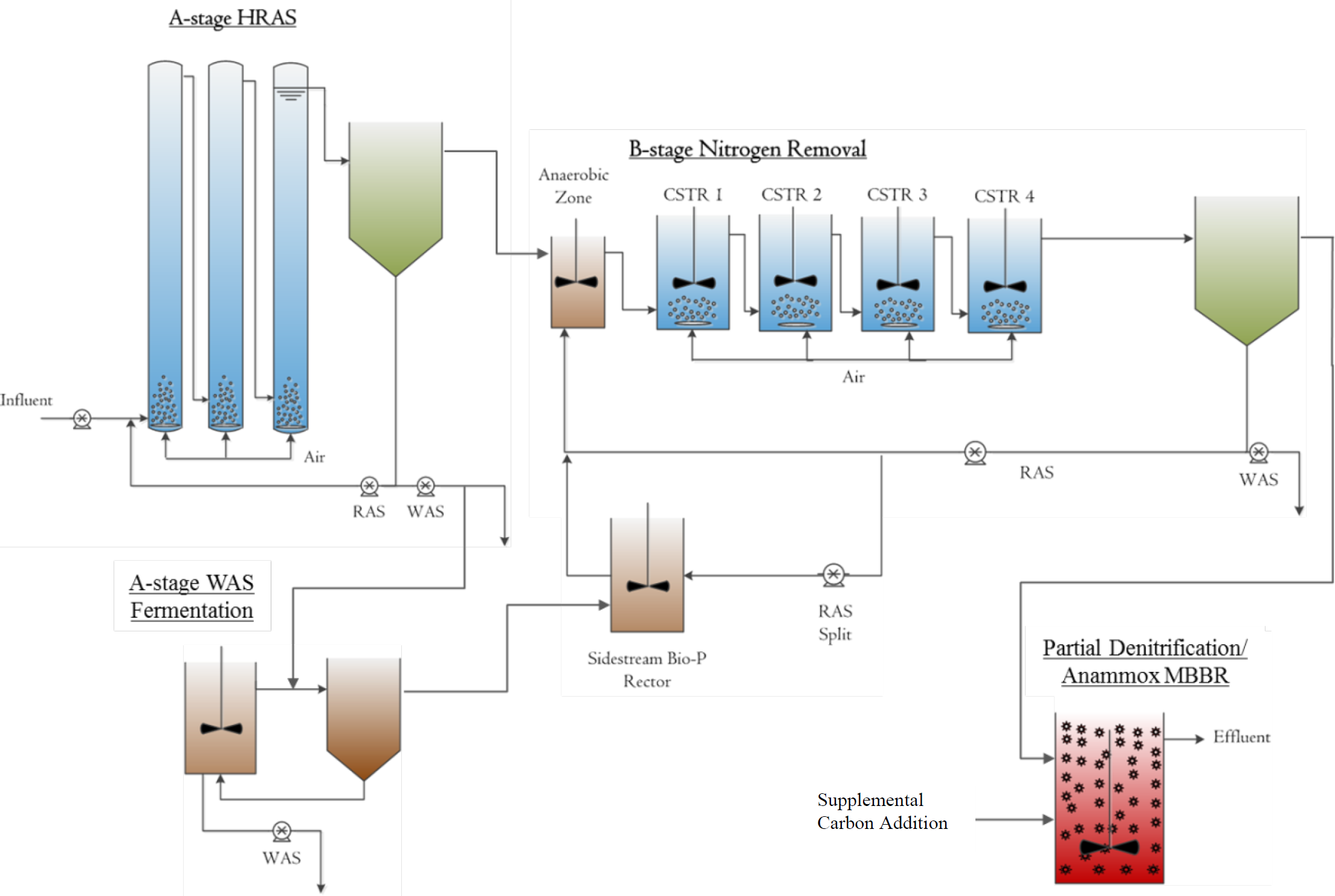
The process-flow schematic of the A/B pilot plant as utilized during the experiment. The process is comprised of the High-Rate Activated Sludge (HRAS) A-Stage, the thickener/fermenter for A-Stage WAS fermentation, the B-Stage Nitrogen Removal Process with Sidstream Bio-P reactor, and the polishing PdNA MBBR.

The sidestream bio-P reactor (SBPR) HRT varied depending on the percentage of RAS sent to the sidestream. A portion of the A-stage WAS was fed to a combined fermenter and thickener, and the fermentate was added to the SBPR. The volume of the fermenter was 190 Liters and the thickener was 340 Liters. The A-stage WAS was fed into the thickener along with the fermenter effluent, and then the underflow of the thickener was fed to the fermenter. In this way, the influent provided elutriation of the fermenter effluent. The effluent from the thickener (fermentate) flowed into a 15-Liter tank, where it was pumped into the bottom of the SBPR reactor (volume = 150 Liters). The influent structure to the SBPR consisted of a one-foot piece of PVC pipe, where the RAS and the fermentate were mixed prior to entering the reactor. The influent structure to the anaerobic selector also consisted of a one-foot section of PVC pipe in which the influent, the RAS, and the SBPR effluent could combine, before entering the reactor. The SBPR was programmed to be mixed for 10 minutes every 3 hours. The anaerobic selector was continuously mixed and covered with ping pong balls to minimize oxygen transfer.

B-stage sensors included a dissolved oxygen (DO) sensor in CSTR 1, CSTR2, CSTR 3, and CSTR 4 (Insite Model 10, LA, USA), ammonium in CSTR 4 (WTW VARiON, Germany), nitrate and nitrite in CSTR 4 (s::can Spectro::lyser, Austria).There was also a NOx sensor in the MBBR (Hach Nitratax plus sc, Loveland, CO). The sensors were connected to a programmable logic controller (PLC) and then used for control as explained below. The B-stage was intermittently aerated using ammonia versus NOx (AvN) control (Regmi et al., 2014); this was achieved via two independent control loops. The first PID control loop determined how often the system was aerating. All aerobic reactors began and ceased aeration at the same time. Thus the first control variable was the aerobic fraction, i.e. the air-on time (variable) divided by total cycle time (fixed); the process variable in this control loop was the NOx/NH3 ratio in CSTR4 (as measured by the ammonia, nitrite, and nitrate sensors). The second PID control loop controlled the aeration during the air-on times. Each reactor had a motor operated valve (MOV) before the diffusor, and the valve position (control variable) for each reactor was adjusted to maintain a DO concentration (process variable) within that reactor. The DO concentration during aerobic cycles was maintained at 2 mg/L in each and the total cycle time varied from 16 to 20 minutes for the duration of the test. Additionally, the carbon dosing pump feed rate for the PdNA MBBR was varied (via PID controller) to achieve a desired NOx concentration.

### 2.2 Analysis

Performance was monitored through the collection of 24-hour composite samples using automated refrigerated samplers (ISCO 3710 Sampler, Teledyne ISCO, Lincoln, NE). The samplers extracted 250 mL at one-hour intervals allowing average daily influent and effluent characteristics to be measured. Total and volatile suspended solids (TSS and VSS) were captured on 1.5 μm filters and analyzed using Standard Methods 2540D and 2540E respectively (APHA, 2012). Total and soluble COD, OP, total ammonia nitrogen (NH_4_^+^-N + NH_3_-N), NO_2_^−^-N, and NO_3_^−^-N were measured with HACH TNTplus kits (Loveland, Colorado) and a HACH DR2800 spectrophotometer (Loveland, CO). Nutrient and soluble COD samples were filtered through 0.45 μm and 1.5 μm filters respectively. Particulate COD (pCOD) was calculated as the difference between total COD and sCOD (1.5 μm filtered). The sCOD measurement includes readily biodegradable and colloidal COD fractions. Daily pH and temperature readings of the reactors were recorded using a handheld pH and temperature meter (Beckman Coulter, Brea, CA). Daily spot-check readings for DO were recorded using a handheld DO sensor (Insite Model 10, LA, USA).

Concentrations of acetic, propionic, butyric, isobutyric, valeric, isovaleric, and caproic acids were determined by GC-FID as previously described in Kinyua et al. (2017). In short, the A-stage WAS and fermentate samples were filtered through a 0.22 µm filter prior to injection. VFA concentrations were determined from a 0.5-100 mg/L calibration curve prepared from commercial standards (Absolute Standards, Hamden, Connecticut).

#### 2.2.1 PAO Uptake and Release Tests

Eight Liters of mixed liquor were collected in a batch reactor from the last CSTR. The reactor was covered and anaerobic conditions were established. Sodium acetate stock solution (10,000 mgCOD/L) was added to reach 300 mgCOD/L in the reactor. Samples were collected every 15 minutes and measured for NO_3_^−^, NO_2_^−^, OP, and sCOD. After the anaerobic release phase, the sample was then split into two four-Liter reactors, one for aerobic OP uptake, and one for anoxic OP uptake. The aerobic reactor was aerated to maintain a DO concentration between 3 and 4 mg/L. Samples were collected every 15 minutes and analyzed for OP. The anoxic reactor was covered and spiked with potassium nitrate stock solution (10,000 mgN/L) to a concentration of 20 mgN/L in the reactor. Samples were collected every 15 minutes, immediately filtered through 0.45μm cellulose acetate filters, and analyzed for OP, sCOD, NO_3_^−^, and NO_2_^−^.

#### 2.2.2 Residual OP Uptake Test

A 4 L sample of sludge was taken from the last aerated reactor in the process. Temperature was controlled at 20°C via submersion in a water bath. The DO concentration was manually maintained between 2.5 and 4 mg/L using diffused compressed air. The pH was manually maintained at approximately 7.5 through the addition of sodium bicarbonate. The sample was spiked with KH2PO4 to achieve an initial OP concentration of 25 mgOP-P/L. Samples were taken every 15 minutes and at least 5 samples were taken for each test.

#### 2.2.3 AOB and NOB Rate Tests

To measure maximum AOB and NOB activity, a 4 L sample was collected from the 4^th^ CSTR and aerated for 30 minutes to oxidize any remaining COD. The sample was then spiked with NH_4_Cl and NaNO_2_ so that initial concentrations were 20-30 mg NH_4_^+^-N/L and 2-4 mg NO_2_^−^-N/L respectively. Temperature was controlled at 20°C via submersion in a water bath. The DO concentration was manually maintained between 2.5 and 4 mg/L using diffused compressed air. The pH was manually maintained at approximately 7.5 through the addition of sodium bicarbonate. The activity tests were conducted for 1 hour with sample collection every 15 minutes. Samples were analyzed for NH_4_^+^-N, NO_2_^−^-N, and NO_3_^−^-N as described above. The AOB rates were calculated as the slope of the NO_x_-N production and NOB rates were calculated as the slope of the NO_3_^−^-N production. Rate tests were performed every 1-2 weeks.

#### 2.2.4 Long-Term Denitrification Test

An 8 Liter sample was collected from the last CSTR and aerated for 48 hours (DO maintained above 3 mg/L) to deplete all external and internally stored carbon. Then the aeration ceased and the sample was split into two 4 Liter reactors, each covered with a Styrofoam lid. One reactor was to measure the endogenous denitrification rate, and the other to measure the denitrification rate from internally stored carbon, after a period of aeration. For the endogenous rate measurement: once the sample went anoxic, samples were taken every 15 minutes for 3 hours. The rate of NOx reduction was considered the endogenous denitrification rate. For the internally stored carbon test: sodium acetate stock solution was spiked to an initial concentration of 250 mgCOD/L for an anaerobic carbon storage phase. Then the reactor was aerated for one hour. Then the reactor was anoxic again, and samples were collected every 15 minutes for 1.5 hours. The NOx rate was considered the denitrification rate from internally stored carbon, or the post anoxic denitrification rate, since external rbCOD was depleted in the aerated phase. Samples were filtered through 0.45 cellulose acetate filters and analyzed for NO_3_^−^-N, NO_2_^−^-N, NH_3_-N, COD, and OP.

#### 2.2.5 External Carbon Independent Denitrification (ECID) Rate Tests

A 4 L sample was collected from the last CSTR and covered with a Styrofoam lid. Potassium nitrate stock solution was added so that the initial NO_3_-N concentration was 10-20 mg/L, and sodium nitrite stock solution was added so that the initial NO_2_-N concentration was 2-6 mg/L. No external carbon sources were added. Samples were taken every 15 minutes for 1 hour, filtered through 0.45 um cellulose filters, and analyzed for NO_3_^−^-N, NO_2_^−^-N, NH_3_-N, COD, and OP.

## 3 Results and Discussion

### 3.1 Pilot Operating Variables

The pilot process was operated for over 1200 days. During this period the A-Stage operation was held relatively constant; in the B-stage, the mainstream SRT, sidestream SRT, aerobic fraction, RAS-split percentage, and VFA addition were each varied during the experiment to cover a range of operational conditions and determine optimal operational settings for effluent quality and process stability. The A-stage WAS fermenter was operated at different SRTs to optimize VFA production and fractionation, and the MBBR was operated with and without supplemental carbon (acetate, glycerol, or methanol) addition. A summary of the relevant operational ranges for the B-Stage is provided in Table 1. Additional operational data can be found in the supporting information.

**Table 1.** Operational parameters for the B-Stage process. Full range and average values are given for the entire experimental duration. VFA* is the fermentate VFA addition to the sidestream normalized to the forward flow of the process.

| Parameter | Range | Average ± SD | Unit |
| --- | --- | --- | --- |
| Total SRT | 6 – 45 | 15 ± 6 | days |
| Mainstream SRT | 3.5 – 20 | 8 ± 3 | days |
| SBPR SRT | 0.1 – 8 | 0.7 ± 1 | days |
| SBPR HRT | 3.6 – 22.3 | 5.7 ± 4 | hours |
| RAS-split | 5 – 34 | 27 ± 8 | % of RAS flow |
| Aerobic Fraction | 40 – 80 | 59 ± 10 | % of cycle time |
| VFA* addition | 0 – 100 | 23 ± 18 | mgVFA/L |
| Influent OP-P | 1.1 – 5 | 2.5 ± 0.6 | mgP/L |
| Influent sCOD | 70 – 190 | 110 ± 25 | mgCOD/L |
| Influent TKN | 27.9 – 51.50 | 40 ± 4.2 | mgN/L |

### 3.2 Phosphorus Removal

Across the wide range of operational conditions tested, the amount of VFA added from the A-Stage WAS Fermenter to the SBPR was found to be the primary factor determining the success or failure of biological phosphorus removal. While the importance of VFA addition is not surprising given the metabolism of PAOs, the stability of phosphorus removal once a threshold of VFA*/P (the VFA mass added to the sidestream normalized to the B-Stage influent flow (VFA*) per total influent OP-P concentration) was met, even across varied operation conditions, was significant. Figure 2 shows the relationship between VFA addition and effluent OP throughout the experiment. When sufficient VFA is added to the sidestream, P removal is stable: 93% of measured effluent OP was below 1.0 mgP/L when VFA*/P ratio ≥ 10, compared with 36% of measured effluent OP under 1.0 mgP/L when VFA*/P < 10. This threshold VFA addition for good P removal compares favorably to other published results: VFA/P = 15 for various SBR systems (Coats et al., 2017) and VFA/TP = 7 - 10 for full-scale bio-P processes (note this range is given relative to total P, not OP) (WEF, 2011), and is not far from the suggested theoretical minimum for P-limitation in PAO metabolism (VFA/P = 8.3) (Schuler and Jenkins, 2003). P removal also remained stable at higher VFA addition levels. Adding more carbon to the SBPR than is strictly required for phosphorus removal did not cause any degradation in performance or apparent problematic competition from GAO or other carbon storing organisms.

**Figure 2.**
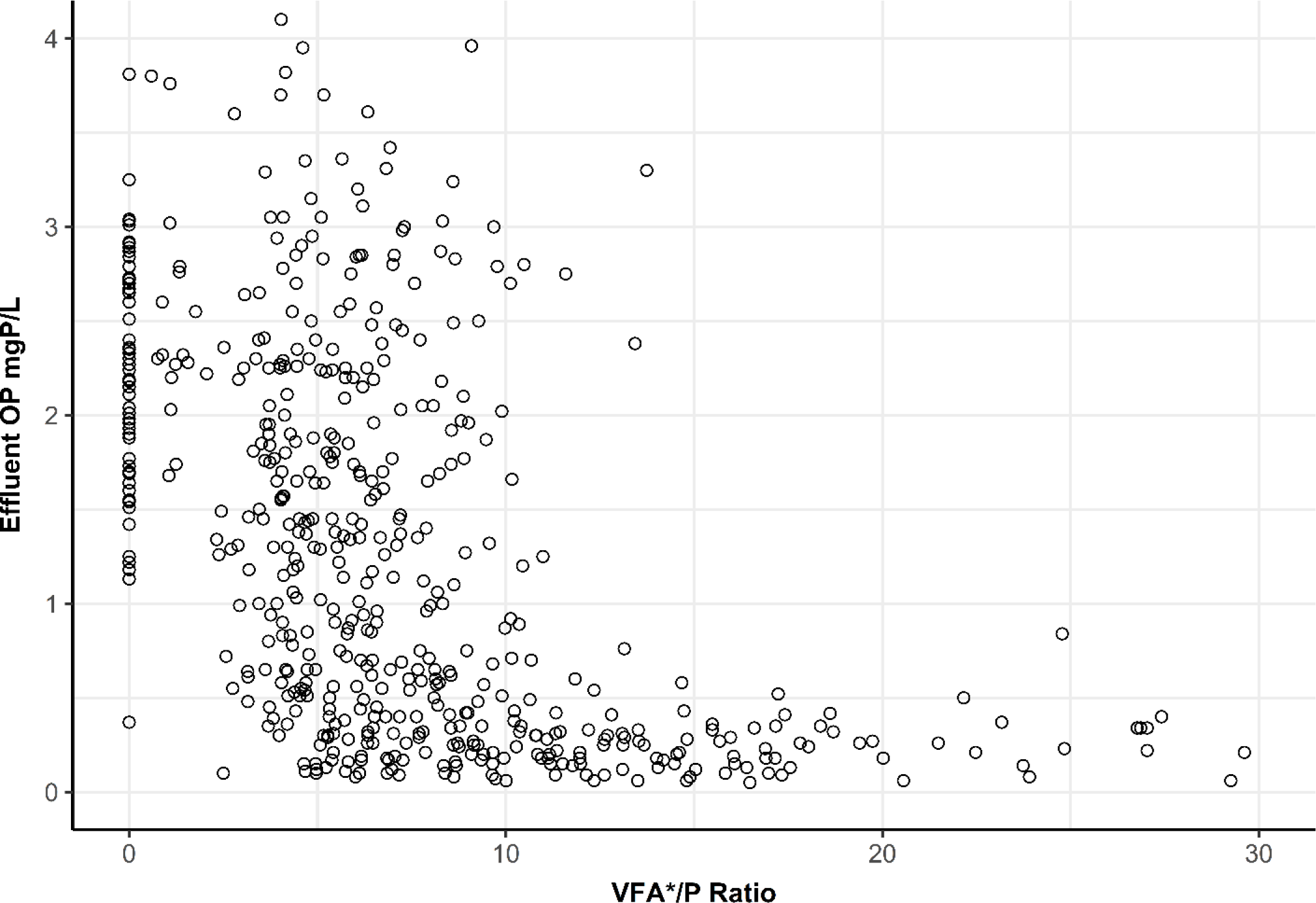
B-Stage effluent OP concentration shown in relation to the amount of VFA added to the SBPR via A-Stage WAS fermentate (here shown in units of VFA*/P ratio: the VFA mass added to the sidestream normalized to the B-Stage influent flow (VFA*) per total influent OP-P concentration).

Figure 2 also shows the effects of zero VFA addition in this system; meaningful biological phosphorus removal could not be achieved with RAS fermentation in the sidestream alone, although this has been observed at many other facilities where carbon is not captured upstream. Bio-P was not successful without VFA addition even when tested at SBPR HRTs over 18 hours and SBPR SRTs above 9 days; this was likely due to the low RAS-Split (less than 10%) required to achieve this longer SBPR HRT.

The PAO uptake/release activity test results correlated well with phosphorus removal performance and with VFA addition, as expected. dPAO were almost never detected during these batch tests. The maximum specific anaerobic P release rate correlated with the process VFA*/P ratio (r = 0.55, p < .01), as was the maximum specific aerobic P uptake rate (r = 0.63, p < .01). The results of these anaerobic P release portion and the aerobic P uptake portion of the tests are shown in relation to the VFA*/P ratio in Figure 3. A marked increase in maximum specific release and uptake rate is seen in both portions of the test until sufficient VFA addition is reached (VFA*/P = 10); after which the rate increases significantly less or even declines slightly (Figure 3). Above this VFA*/P threshold, the maximum specific anaerobic P release rate was higher than in other studies of sidestream bio-P systems (2.85 – 5.3 mgOP-P/gVSS/h) but within ranges found in conventional bio-P processes (3.2 – 31.9 mgOP-P/gVSS/h) (Onnis-Hayden et al., 2020). The maximum specific aerobic P uptake rate was similarly higher than found in studies of sidestream bio-P systems (0.6 – 2.6 mgOP-P/gVSS/h) and at the upper end of the range of conventional bio-P processes (0.7 – 7.6 mgOP-P-gVSS/h) (Onnis-Hayden et al., 2020). This relationship demonstrates the PAOs in the system at the time of the test being carbon-limited up to the point of sufficient VFA addition and phosphorus-limited thereafter (little increase in maximum rate with additional VFA addition). It should be noted the VFA*/P ratio indicates the VFA addition to the process at the time the test was performed, not the VFA addition to the P uptake and release test; in the bio-P activity test VFA is always added in non-limiting amounts to achieve the maximum rate.

**Figure 3.**
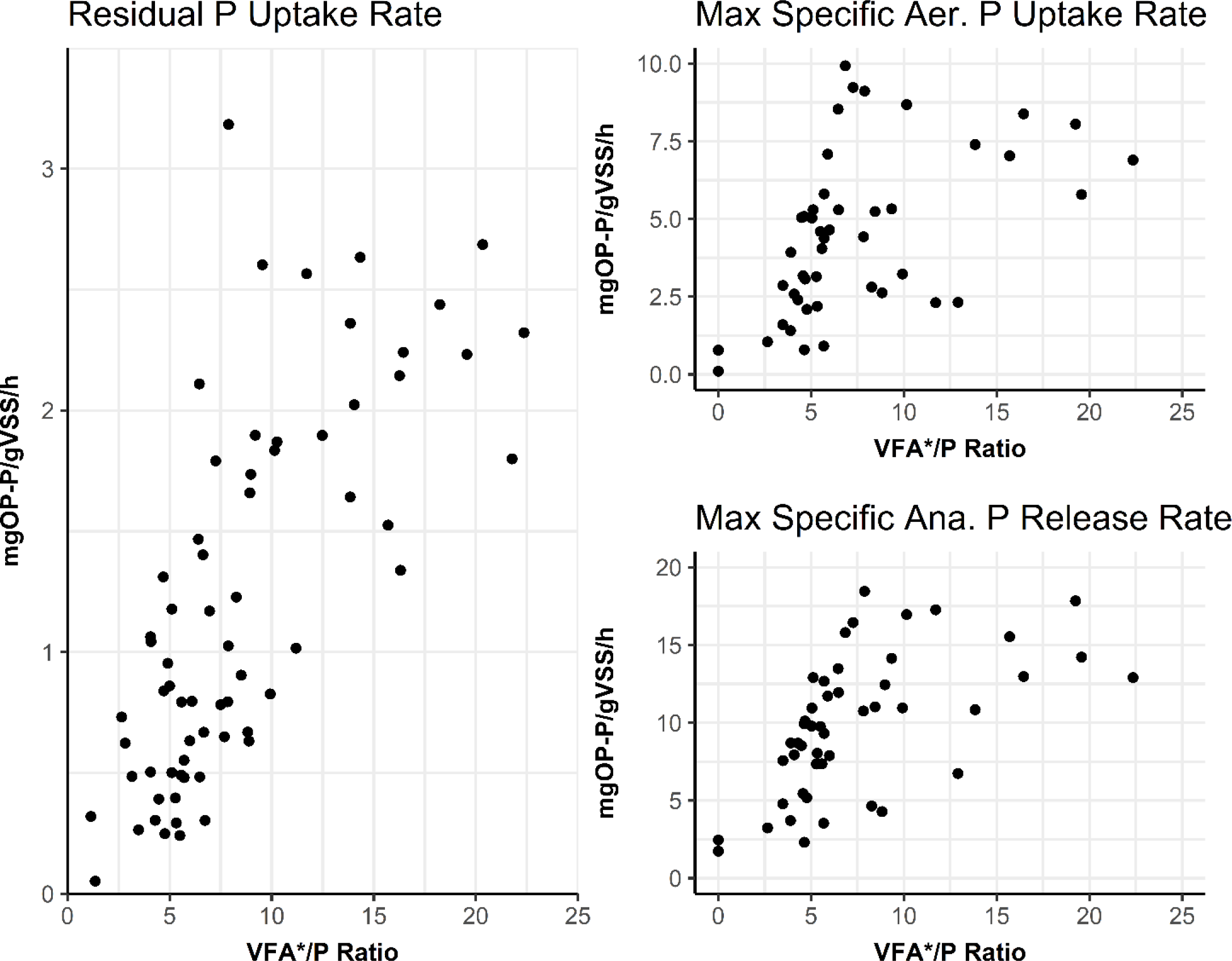
Specific P uptake and P release rates from the PAO maximum activity test (Max Aerobic P Uptake Rate and Max Anaerobic P Release Rate) and the RUTOP test (Residual Uptake Rate) shown in relation to the amount of VFA added to the SBPR via A-Stage WAS fermentate at the time the batch test was performed (here shown in units of VFA*/P ratio: the VFA mass added to the sidestream normalized to the B-Stage influent flow (VFA*) per total influent OP-P concentration).

The results of the Residual OP Uptake (RUTOP) test are also shown in Figure 3. This test measures the residual or remaining activity of PAOs at the end of the aerobic process via additional aeration and as such the rates are much lower than the maximum rates. A similar relationship is seen between the maximum release/uptake rates and the residual rates at VFA*/P levels below 10, showing that at insufficient VFA addition levels the PAOs were likely carbon-limited. The RUTOP rate increased with elevated VFA*/P ratio up until 25 beyond the theorized required value of 10, whereas the maximum P uptake rates plateaued. When sufficient VFA*/P is achieved, the bio-P activity became phosphorus-limited as the effluent P concentration is low. The continuous increase in RUTOP rate, but not maximum uptake rate, beyond this threshold level of VFA*/P suggests that the PAOs are able to utilize the VFAs and store sufficient intracellular carbon storage to enable the continuous P uptake at sub-maximum rates, despite being phosphorus limited. Finally, the RUTOP rate test is also a far-less time consuming test than the maximum activity tests, and because it correlates well with process performance (and carbon addition), may be a useful measure operationally.

#### 3.2.1 Impact of BNR SRT, sidestream SRT, and other operational parameters on bio-P performance

This relationship between VFA*/P and successful phosphorus removal held across the range of B-Stage SRTs (3.5 – 20 days) tested during the experiment. The total B-Stage SRT was not significantly correlated with effluent phosphorus concentration (r = -.59, p = .21). The SRT of the SBPR had a very small but statistically significant correlation with effluent phosphorus concentrations (r = 0.11, p = .014), but when the data were split by VFA addition, this correlation did not remain significant in either the sufficient VFA addition group (VFA*/P > 10, r = .13, p = .22) or the insufficient VFA addition group (VFA*/P < 10, r = .1, p = .08). The lack of correlation between the effluent phosphorus and the total SRT or SBPR SRT could indicate SRT-dependent processes like hydrolysis and fermentation were not significant contributors to PAO activity. This lack of correlation indicates this process is relatively robust, as effluent quality was not diminished by low (under 6 days) or high (over 20 days) SRT, although sufficient VFA must be supplied.

Another significant factor in the operation of sidestream biological phosphorus removal is the RAS split percentage, or the fraction of RAS that is diverted to the sidestream instead of being returned to the mainstream portion of the process. This also impacts the HRT of the sidestream, assuming the volume is held constant. In this experiment, the RAS split percentage was primarily held constant at 30%. Starting at day 483, for 90 consecutive days, tests were conducted by lowering the RAS split percentage (as low as 5%) to increase the HRT of the SBPR. At the same time VFA addition was reduced below the sufficient threshold and eventually stopped, to determine if internal hydrolysis and fermentation was sufficient to sustain phosphorus removal. These tests were not successful, as the effluent phosphorus was observed under 1.0 mgP/L only 6% of the time. The RAS split percentage was then returned to 30% and VFA addition restarted. It was not possible to tell in this test if the limiting factor for bio-P was hydrolysis/fermentation, insufficient exposure of the PAO biomass to sidestream conditions, or a combination of the two. Larger sidestream reactors could be tested in this configuration, but this may also be a limitation of the carbon-limited B-Stage process that conventional sidestream bio-P processes do not need to contend with.

Numerous other operational factors had little impact on the effectiveness of phosphorus removal in the system. The SBPR was intermittently mixed, and this mixing was varied from continuous mixing to mixing for only 20 minutes per day, and was most frequently set at 24 minutes per day, or 1.67% of the day (8 mixing events per day, 3 minutes each). Varying the mixing allowed more or less solids to accumulate in the SBPR and varied the depth and concentration of the solids in the sludge blanket that developed. This may have allowed for increased fermentation and hydrolysis within the SBPR, but if this did occur, it was not a significant contributor to the phosphorus removal as the impacts of SBPR mixing on effluent P concentrations were very small, though statistically significant (r = .07, p = .04). The Aerobic Fraction of the process also had a very small but statistically significant correlation with effluent P concentration (r = -.08, p = .03), demonstrating that the PAOs in the system were capable of aerobic phosphorus uptake across the range of Aerobic Fractions tested (40-80% aerobic time), and did not require continuous aeration, as long as sufficient VFA was added to the system. The A-stage effluent sCOD/P ratio had a statistically significant but very low correlation with effluent P concentrations (r = -.11, p = .01). The speciation of the VFA produced by the A-Stage WAS Fermenter varied during the experiment, but was predominantly comprised of acetate and propionate. The ratio of acetate to propionate in the fermentate varied from 0.6 – 3.1 with a median value of 1.1, but did not significantly correlate to effluent P concentrations (r = 0.13, p = 0.14).

### 3.3 Nitrogen Removal

The impacts of incorporating sidestream bio-P into the A-B process were not limited to phosphorus removal, nitrogen removal was affected as well. Nitrification and denitrification were performed alternatingly within the B-Stage through the cycling of aerobic and anoxic periods to produce an effluent that contained a ratio of ammonia and nitrate+nitrite that would be conducive to downstream treatment via anammox in the MBBR. The sidestream process and VFA addition impacted the nitrogen removal performance through two routes. First, TIN removal via denitrification was enhanced due increased carbon loading to the system via the addition of WAS fermentate; some of this carbon was also internally stored and contributed to post-anoxic denitrification. Second, the downstream anammox process was enhanced through the generation of nitrite in the B-Stage through partial-denitrification as well as NOB out-selection during different phases of the pilot. This nitrite then contributed to PNA in the downstream anammox MBBR.

#### 3.3.1 TIN Removal via Denitrification

Overall, increased fermentate addition (higher VFA*/P ratio) lead to increased TIN removal in the B-Stage as shown in Figure 4. This increased TIN removal could be attributed to overall increased carbon loading leading to more denitrification during the anoxic cycles, or increased intracellular storage of carbon that was used for denitrification, or a combination of the two.

**Figure 4.**
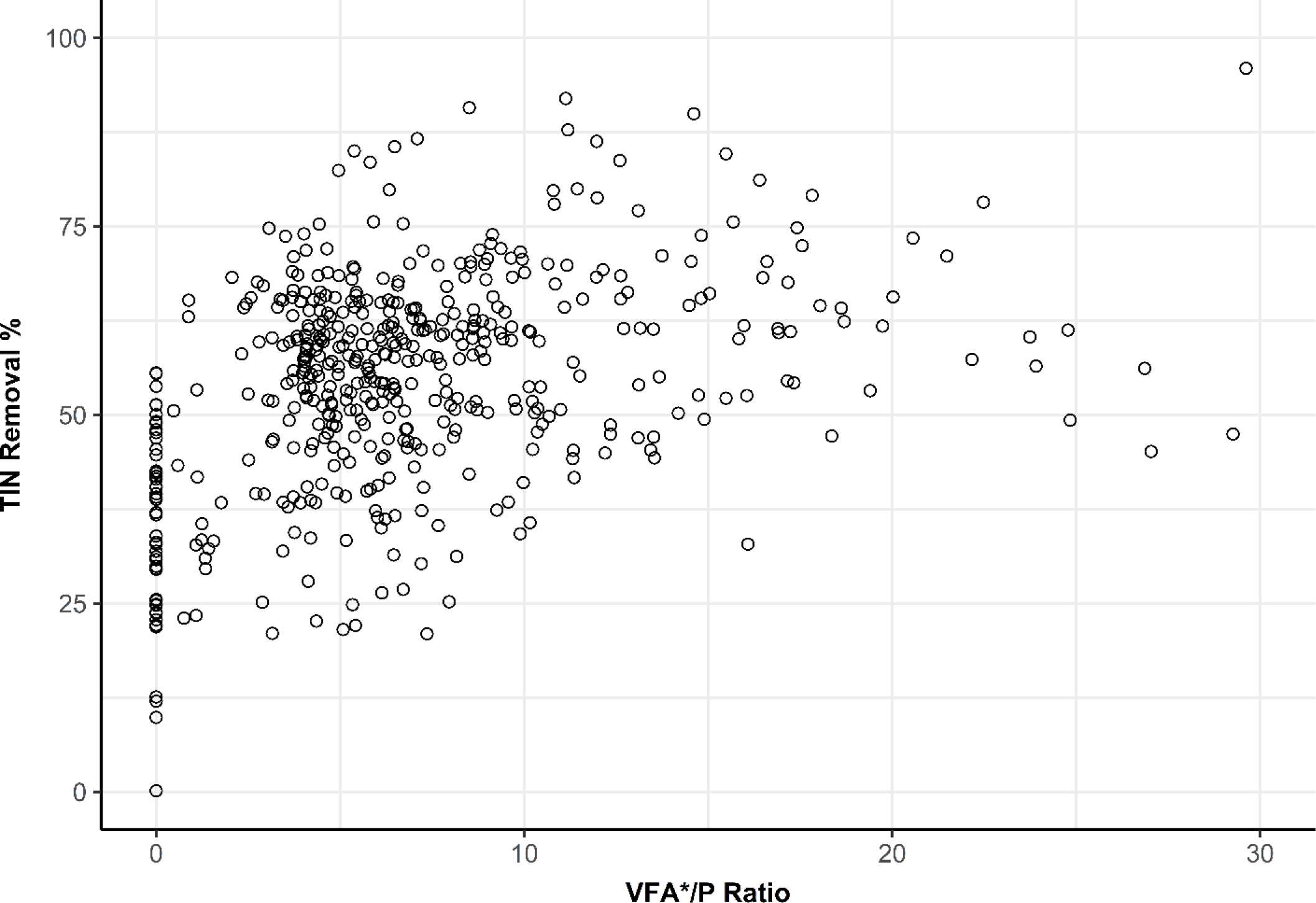
Overall B-Stage TIN removal percentage as a function of carbon addition to the sidestream (given in VFA*/P Ratio units). TIN removal shown is only for the B-Stage (intermittently aerated) portion of the process, and does not include additional TIN removal in the downstream anammox MBBR.

To better understand the phenomena responsible for the increased TIN removal, denitrification rate tests were conducted with mixed liquor sampled from the last reactor in the B-Stage process without addition of exogenous carbon (the ECID batch test). These samples thus had very little or no rbCOD remaining, and the primary drivers of the TIN removal rate would be intracellularly stored carbon and endogenous denitrification. The results of these batch tests are shown in Figure 5. When no fermentate is added to the sidestream (VFA*/P = 0) it can be assumed that the specific denitrification rate is determined primarily by endogenous decay. As more COD is added to the sidestream, the specific denitrification rate increases; because the specific rate takes the VSS of the mixed liquor into account, this cannot merely be due to increased carbon loading causing increased heterotrophic growth and higher endogenous rates, and is likely due to some amount of carbon from the sidestream being stored and still available at the end of the aerated process for further nitrogen reduction. Additional batch testing was done to further explore these mechanisms.

**Figure 5.**
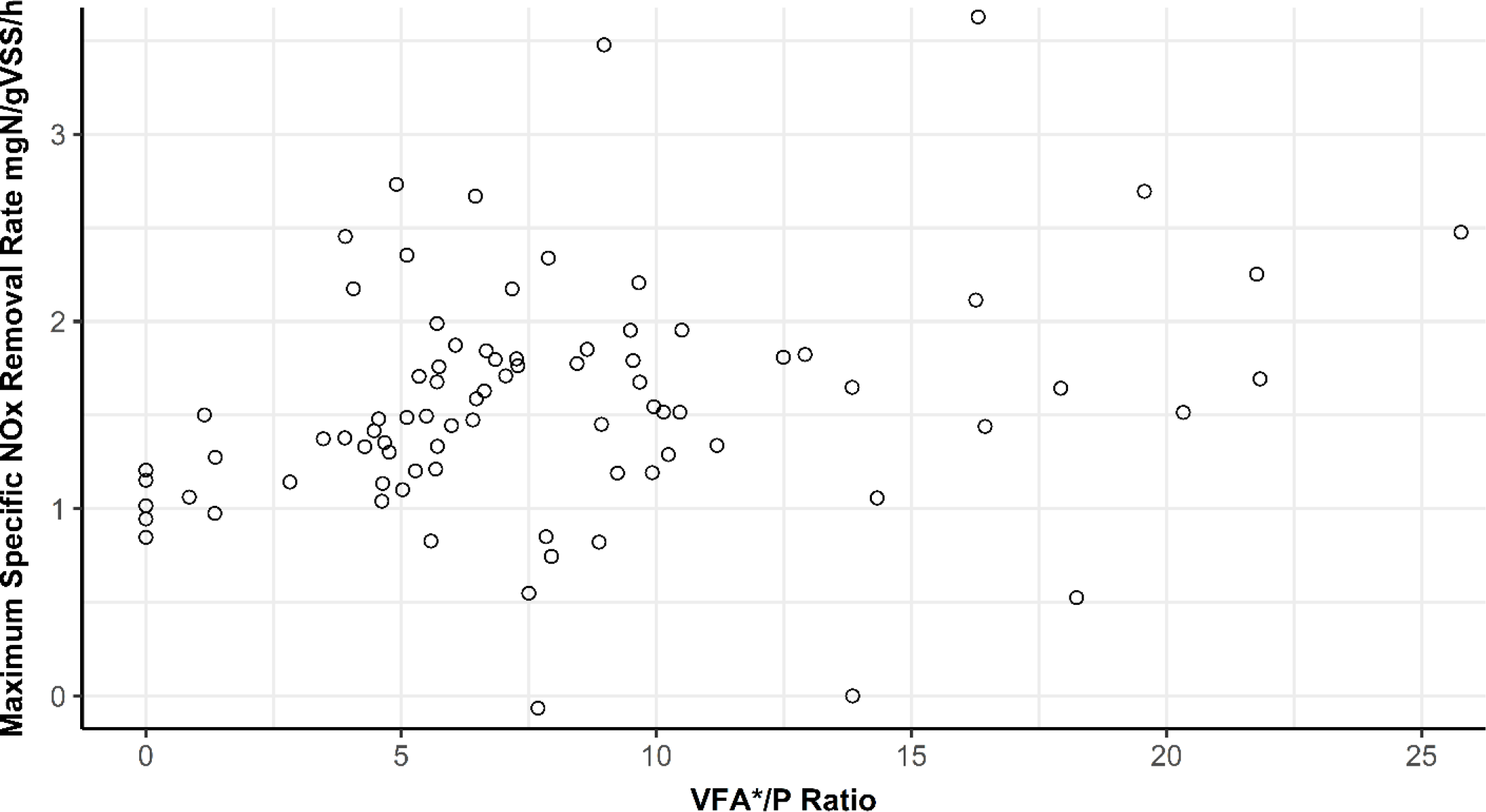
External carbon independent denitrification (ECID) batch test NOx removal rates shown as a function of carbon addition to the sidestream (in units of VFA*/P ratio). Specific denitrification rates increased with increased carbon loading to the sidestream.

#### 3.3.2 Post-anoxic denitrification

The term post-anoxic denitrification refers to an anoxic zone that follows an aerated zone, without external carbon addition. This is equivalent to the anoxic periods of intermittent aeration, once external carbon sources have been depleted. Post-anoxic denitrification typically implies endogenous denitrification, but it is not always specified whether this includes denitrification from internally stored carbon, as internal storage compounds are not typically measured. The long-term denitrification test was performed on Day 651 in order to separately measure the endogenous and post-anoxic denitrification rate (from acetate storage only) (Figure 4). There was no nitrite accumulation in this test, so the following denitrification rates are the rate of nitrate reduction. After the sample was aerated for 48 hours, the measured endogenous denitrification rate was 0.48 mgN/gVSS/hr. Typical endogenous denitrification rates are in the range of 0.2-0.8 mgN/gMLSS/hr (Kujawa and Klapwijk, 1999).

**Figure 6.**
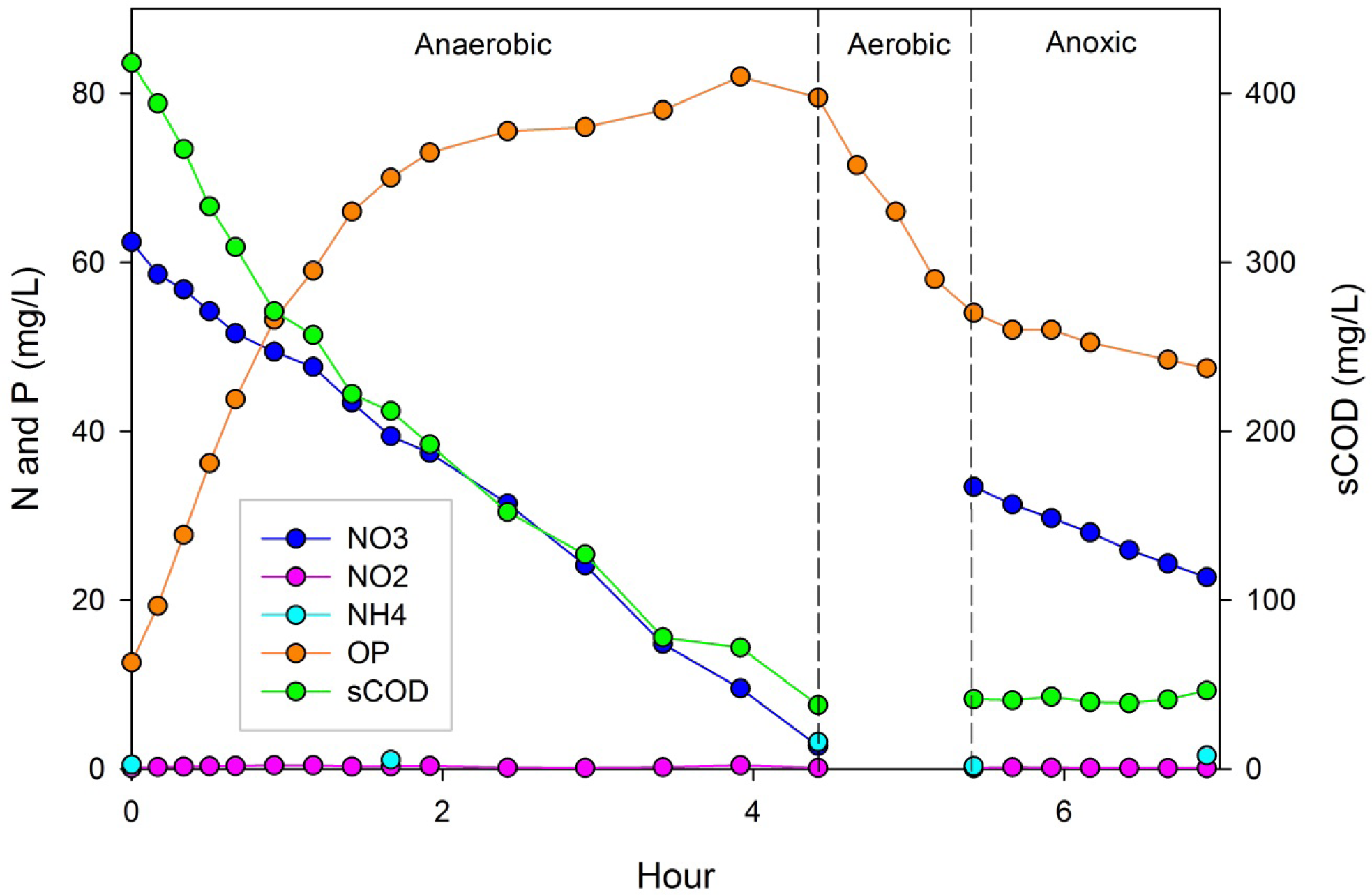
Results from the long-term denitrification test. The first period is labeled “anaerobic” even though there is nitrate present because this is the OP release and VFA storage period, followed by the aerobic period, then anoxic.

The rate of denitrification with acetate as the only carbon source was 2.4 mgN/gVSS/hr. There may have been some endogenous denitrification occurring during the anoxic period with acetate, but it can be assumed that there were no internal carbon stores at the start of the anoxic period after 48 hours of aeration. The acetate was first utilized both for OP release by PAO and for denitrification by OHO (hour 0 to 2), and then just for denitrification once OP release ended (hour 2 to 4). After a period of 1 hour of aeration to deplete any residual rbCOD, the post-anoxic denitrification rate was 1.3 mgN/gVSS/hr which is less than the rate with acetate as the carbon source, but larger than the endogenous denitrification rate. Since the only available carbon source after 48 hours of aeration was the spiked acetate, which was then depleted, it can be assumed that internally stored carbon is the electron donor for denitrification. In this case, there was marginal OP uptake in the last anoxic period (0.76 mgP/gVSS/hr), but denitrification in the ECID tests was not typically associated with OP uptake. The average NOx reduction rate in the ECID batch tests during this time (Day 651 to Day 689) was 1.4±0.2 mgN/gVSS/hr. Interestingly, this is close to the post-anoxic rate that was observed in the batch test after acetate storage (1.3 mgN/gVSS/hr). Vocks (2008) performed similar batch tests using mixed liquor from three full-scale facilities: two UCT processes (with bio-P), and one pre-denitrification process without bio-P. The average post-anoxic denitrification rates for the UCT processes were 1.6 mgN/gVSS/hr (n=10) and 1.4 mgN/gVSS/hr (n=2), respectively, with values ranging from 0.7-2.7 mgN/gVSS/hr and 1.2-1.5 mgN/gVSS/hr. For the facility without bio-P the average was 0.6 mgN/gVSS/hr (n=3) ranging from 0.4-0.7 mgN/gVSS/hr. It is possible that post-anoxic denitrification may be occurring more frequently than is recognized in combined bio-P and N removal processes, because post-anoxic denitrification without external carbon addition in full scale processes is not common.

Although the concept of denitrification from internal carbon storage is not new (Alleman and Irvine, 1980; van Loosdrecht et al., 1997), there has been a recent interest in exploiting post-anoxic denitrification (and potentially nitrite accumulation) combined with bio-P (Winkler et al., 2011; Chen et al., 2013; Liu et al., 2017; Zhao et al, 2019; X. Wang et al, 2019; Bauhs et al., 2022). These studies propose utilizing variations of an anaerobic/aerobic/anoxic configuration. Winkler et al. (2011) reported greater than 99% inorganic nitrogen and phosphorus removal, and although GAO were thought to be responsible in part for the post-anoxic denitrification, bio-P performance was not compromised. Liu et al. (2017) and Zhao et al. (2019) proposed nitrite accumulation via partial nitrification, followed by anoxic denitrification of the nitrite utilizing internally stored carbon. They recognized the challenge of maintaining post-anoxic denitrification by GAO, because of the competition with PAO for the VFA. X. Wang et al. (2019) operated two SBRs in series, the first was operated in anaerobic/anoxic/aerobic phases (with added nitrate in the influent), and nitrite was accumulated during the anoxic phase by GAO, followed by an anoxic anammox SBR. Bauhs et al. (2022) identified full-scale BNR facilities that required far less external COD addition than expected, given their nitrogen removal performance, and performed batch experiments on sludge from these facilities. In plants where bio-P was performed alongside nitrification/denitrification, the biomass was capable of significant post-anoxic denitrification via internally stored carbon; at non-nitrifying bio-P plants this was not the case. These denitrification rates correlated positively with anaerobic zone VFA loading and with aerobic phosphorus uptake rates, but anoxic phosphorus uptake (dPAO activity) was not found. Overall, it appears that utilizing internally stored carbon for denitrification in bio-P processes is promising, especially if it results in nitrite accumulation, as long as the PAO are not outcompeted. It would be preferable if the denitrification was being performed by PAO, but it is still unclear how to design a process to guarantee dPAO activity.

#### 3.3.3 Nitrogen Removal via Anammox

Nitrogen removal was accomplished in the B-Stage process as well as the downstream anammox MBBR. The B-Stage AvN controller ensured a proportional mix of nitrate/nitrite and ammonia was available in the MBBR influent. When the NOx portion of the B-Stage effluent was primarily comprised of nitrate, supplemental carbon was added to the MBBR to facilitate partial denitrification and thus accomplish PdNA (Klaus et al., 2023). This process worked well and could achieve TIN removal above 80% with significantly less carbon addition than conventional processes. Additionally, supplemental carbon addition was not necessary (or only required in very small amounts) if the NOx portion of the B-Stage effluent was composed primarily of nitrite. This metric was monitored as the Nitrate Accumulation Ratio (NAR), calculated as the ratio of nitrite to NOx in the B-Stage effluent (Figure 7). Significant NAR was achieved through two mechanisms: 1) partial denitrification within the B-Stage via carbon-storing heterotrophs and 2) NOB-outselection and partial nitritation. High NAR allowed the anammox MBBR to operate in PNA mode (instead of PdNA), and to achieve TIN removal above 80% with no supplemental carbon addition.

**Figure 7.**
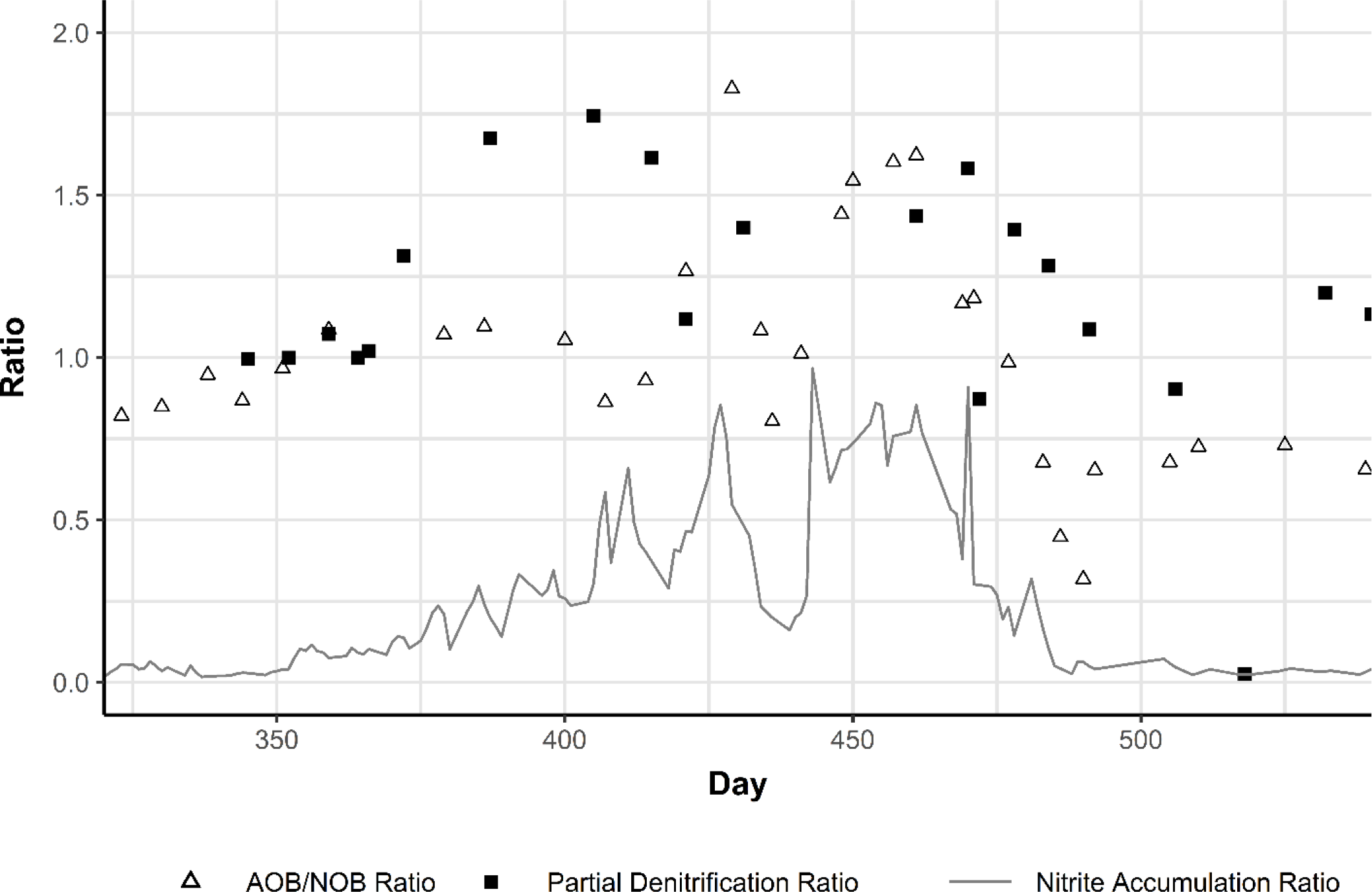
Batch test results and nitrite accumulation in the B-Stage process for the experimental phase. The AOB/NOB ratio is obtained from the weekly maximum nitrification activity batch tests. The Partial Denitrification Ratio is obtained from the weekly ECID tests where maximum denitrification rates are taken with no exogenous COD addition. The Nitrite Accumulation Ratio is the ratio of B-Stage effluent nitrite to effluent NOx.

##### 3.3.3.1 Partial Denitrification via Carbon Storage

The first period of high NAR occurred in the process beginning around day 350. At this point nitrite began to accumulate in the B-Stage effluent and the results of the ECID activity tests began to show a higher denitratation rate relative to the overall denitrification rate. The ratio of these rates is termed the Partial Denitrification Ratio and is shown during this period in Figure 7. The Partial Denitrification Ratio indicates how much nitrite accumulates during these batch tests (where no rbCOD is available and no external COD is added), indicating the preferential use of intracellularly stored carbon for partial versus full denitrification. When the Partial Denitrification Ratio is 1, no nitrite accumulation occurs during the test and the denitratation and denitritation rates are equal. As shown in Figure 7, at the start of this period the Partial Denitrification Ratio was very close to 1.0, and both it and the NAR increased over time until reverting back to 1.0 and below towards the end of the period. During this period the ratio of AOB and NOB maximum activity did not increase much above parity until day 420. To confirm these results a profile of the last B-Stage reactor was taken on day 397, the results of which are shown in Figure 8. The results of this profile demonstrate the nitrite is being generated primarily by partial denitrification: nitrite was consumed (nitratation) during aerobic periods of the intermittent aeration cycle, and produced during the anoxic periods of the cycle (partial denitrification). The nitrate results follow the same but inverse pattern. During this period maximum PAO activity tests were performed which included an aerobic and anoxic P uptake phase; these tests did not show dPAO activity, demonstrating that this NAR was not due to dPAO in the process.

**Figure 8.**
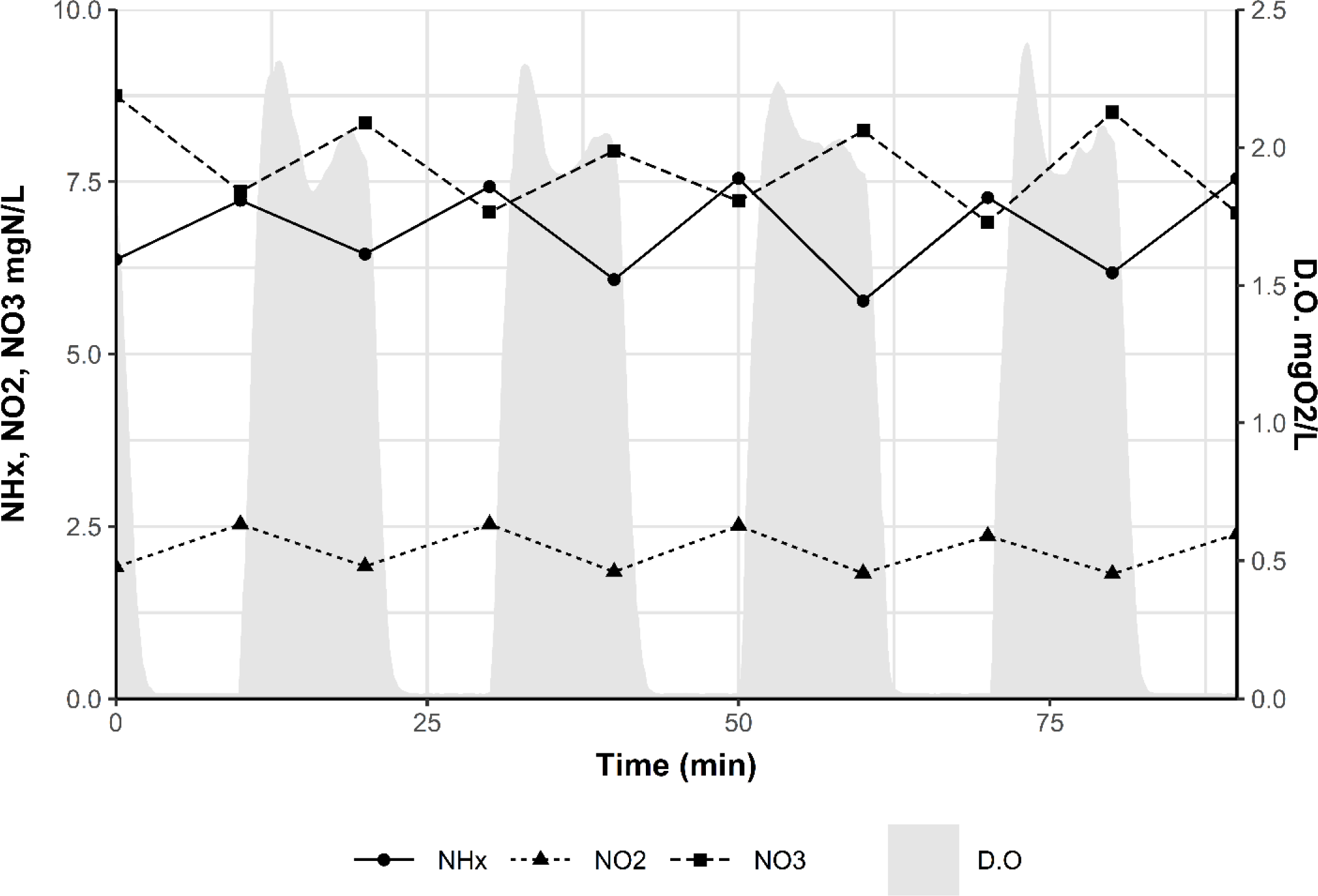
Profile over 90 minutes in the last aerated B-Stage reactor from day 397. Grab samples were taken as the reactor cycled between aerobic and anoxic phases. The nitrite can be seen to accumulate during the anoxic cycle (partial denitrification) and decrease during the aerobic cycle (nitratation).

The results of these batch tests and the profile indicate that the increase in NAR in this period was due primarily to partial denitrification via internal carbon storage, although it appears after the NAR increased the AOB/NOB ratio also increased indicating that NOB out-selection may have played some role as well. To test the resilience of the partial denitrification and determine if it could be sustained only through fermentation and storage within the sidestream reactor, the VFA addition was decreased starting on day 478 and reduced to zero on day 510. This led to a drop in the Partial Denitrification Ratio and ultimately a decrease of NAR down to almost zero.

Figure 9 documents the nitrogen removal performance of the process during this period. At day 430 a temporary drop in NAR and nitrogen removal occurred; this was due to inhibition of nitrifiers from constituents in the raw influent (likely cyanide). The recycle flows responsible for this inhibition were diverted away from the pilot and the nitrogen removal and NAR efficiency resumed shortly thereafter. For a majority of this phase supplemental COD was supplied to the MBBR for PdNA, but this was turned off when the NAR went above 70% to operate in PNA mode. The process was able to perform significant N removal: more than 60% in the B-Stage through nitrification/denitrification and an additional 20-30% removal in the MBBR via anammox.

**Figure 9.**
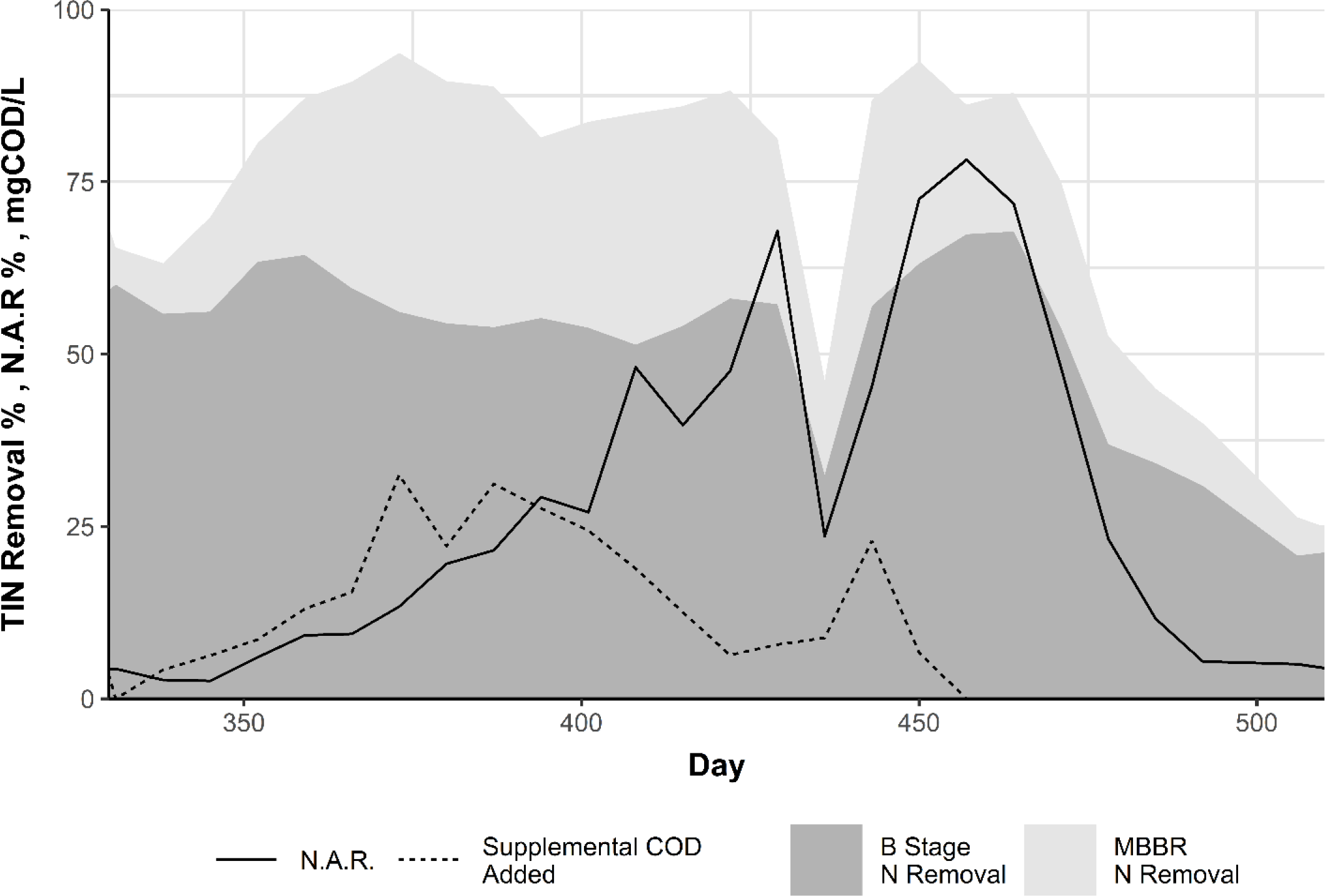
Nitrite accumulation and nitrogen removal performance during the experimental phase. Stacked shaded areas show the percentage of nitrogen removal in the B-Stage and anammox MBBR, total nitrogen removal is the sum of the two areas. Supplemental COD is the concentration of COD added to the anammox MBBR for PdNA (the concentration value shown is normalized to the forward flow of the process).

##### 3.3.3.2 Partial Nitritation via NOB Out-Selection

A second period of high NAR occurred starting around day 750 and lasted approximately 200 days. The results from the nitrification and ECID activity tests are shown in Figure 10 alongside the NAR. Throughout this period the partial denitrification ratio was consistently around 1 indicating that no partial denitrification was occurring. However, the AOB/NOB activity ratio increased drastically as NAR increased, suggesting very high levels of NOB out-selection. A profile taken from the last aerated reactor in the B-Stage confirmed the results, the data of which are shown in Figure 11. There was not an apparent singular cause for the onset of NOB out-selection, but the B-Stage mainstream SRT was lowered at the beginning of this phase to an average of 5.9 days. Aggressive SRT and intermittent aeration is a mechanism for promoting NOB out-selection (Regmi et al., 2014), but as is often the case the effects were transient. Once the NOB out-selection stopped, it was not achieved again even at similar SRTs. VFA addition was varied throughout this period, but it was within typical ranges during the onset of the NOB out-selection.

**Figure 10.**
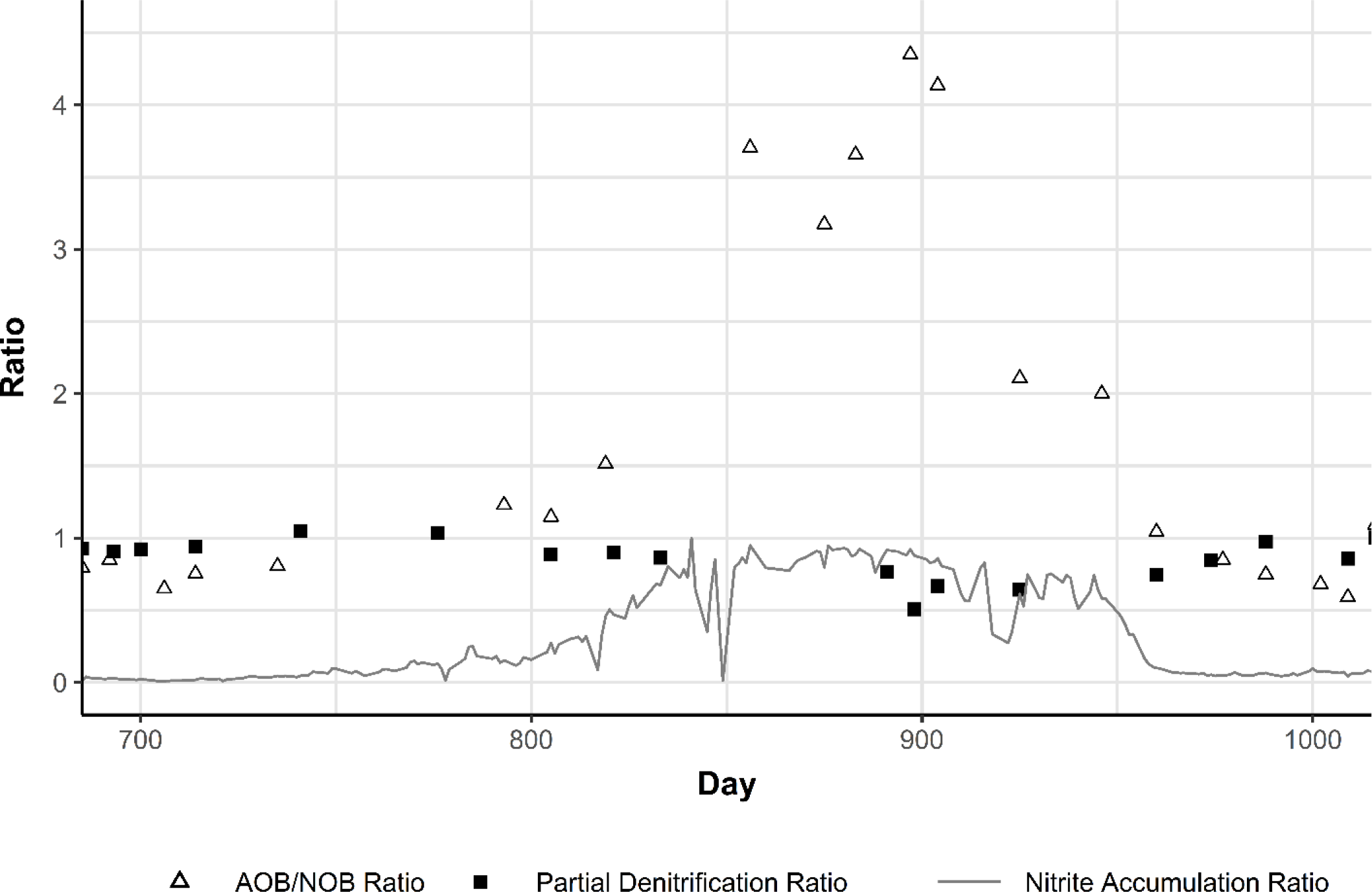
Batch test results and nitrite accumulation in the B-Stage process for the experimental phase. The AOB/NOB ratio is obtained from the weekly maximum nitrification activity batch tests. The Partial Denitrification Ratio is obtained from the weekly ECID tests where maximum denitrification rates are taken with no exogenous COD addition. The Nitrite Accumulation Ratio is the ratio of B-Stage effluent nitrite to effluent NOx.

**Figure 11.**
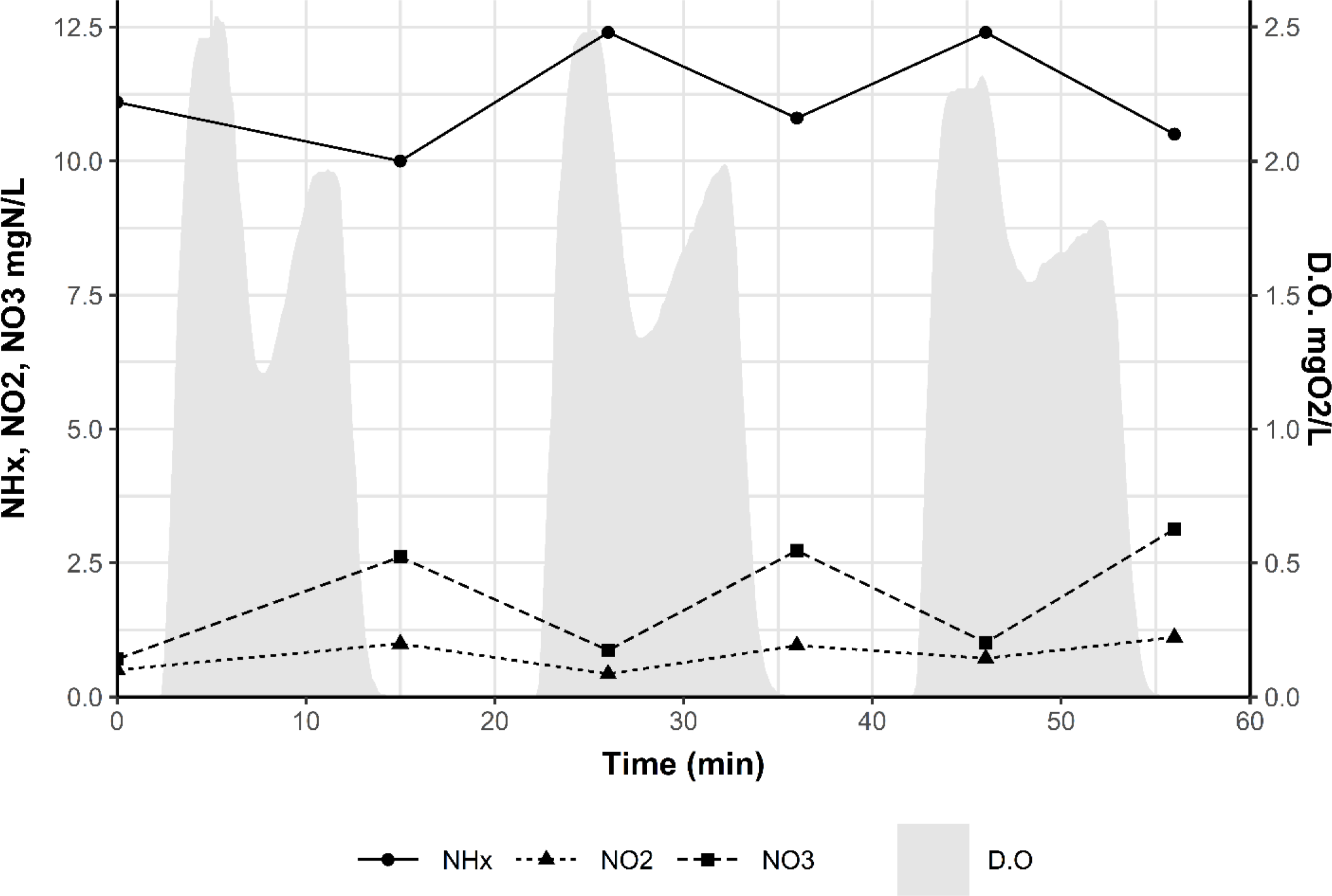
Profile over 60 minutes in the last aerated B-Stage reactor from day 811. Grab samples were taken as the reactor cycled between aerobic and anoxic phases. The nitrite can be seen to accumulate during the aerobic cycle (partial nitritation) and decrease during the anoxic cycle (denitrification).

Figure 12 documents the nitrogen removal performance of the process during this period. There was a brief period where supplemental COD was added to the MBBR for PdNA, but this was turned off. The process was able to perform significant N removal: more than 70% in the B-Stage through nitrification/denitrification and an additional 20% removal in the MBBR via anammox.

**Figure 12.**
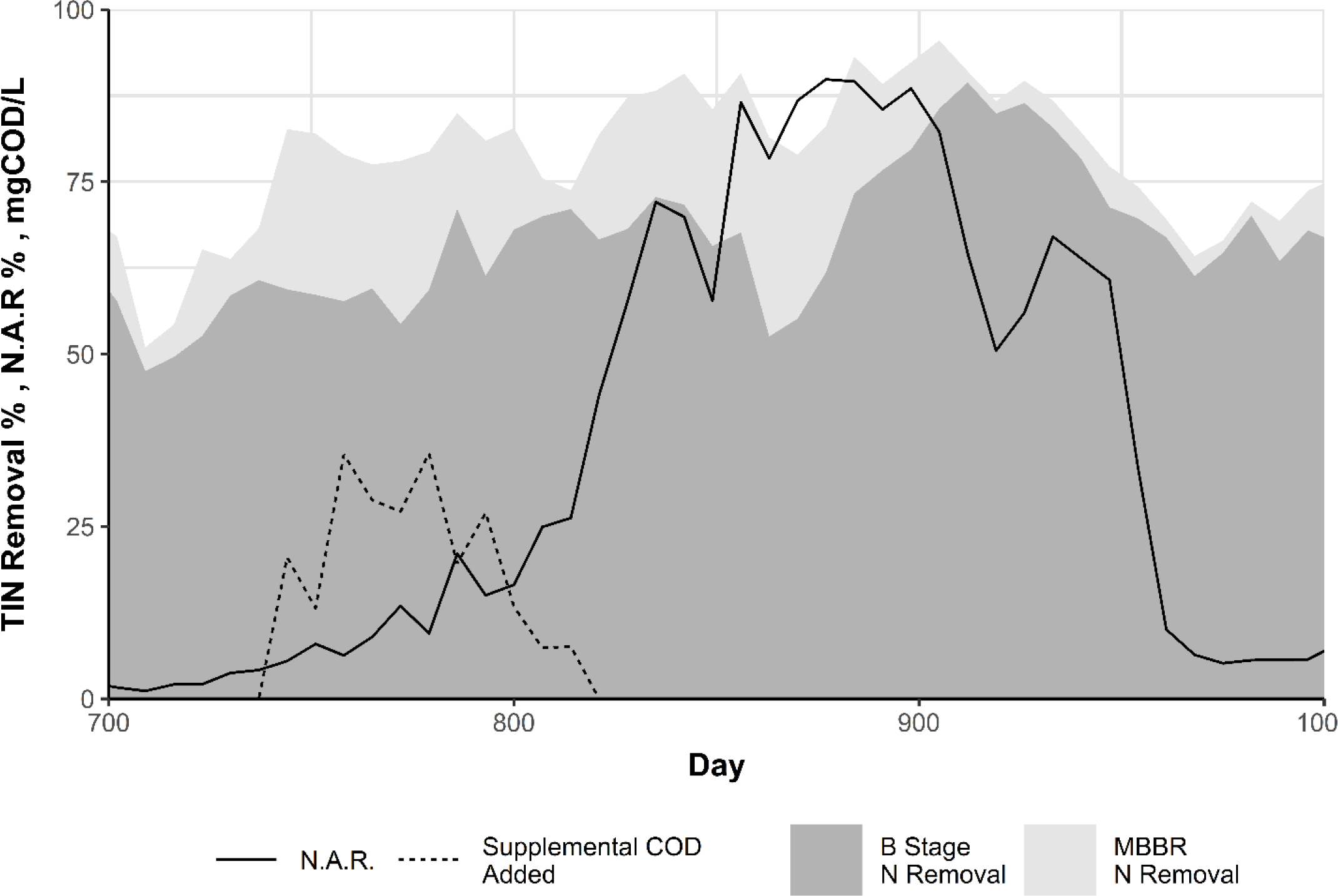
Nitrite accumulation and nitrogen removal performance during the experimental phase. Stacked shaded areas show the percentage of nitrogen removal in the B-Stage and anammox MBBR, total nitrogen removal is the sum of the two areas. Supplemental COD is the concentration of COD added to the anammox MBBR for PdNA (the concentration value shown is normalized to the forward flow of the process).

## 4 Conclusions

A novel process configuration via pilot testing that sought to combine the benefits of carbon diversion and shortcut nitrogen removal (A/B Process), mainstream anammox (MBBR), and sidestream bio-P (with A-Stage WAS fermentate addition) was investigated. The pilot was successful in demonstrating stable biological P removal and low effluent TIN. This system demonstrates that carbon redirection is compatible with biological phosphorus removal, and may be preferable to conventional BNR systems as carbon can be partially removed, transformed into a more desirable form (VFA), and added to the system in only the amounts required to enhance N and P removal.

The key findings were:

- Bio-P with low or no influent VFA was possible and robust as long as sufficient VFA was added to the sidestream system in the form of A-Stage WAS fermentate.
- Bio-P was not inhibited by intermittent aeration and was possible over a wide range of system SRTs.
- Bio-P required a VFA/P ratio of at least 10 gVFA/gP to be added to the sidestream to be successful. VFA/P ratios above this (tested up to 30) did not degrade effluent OP quality.
- VFA fed to the sidestream reactor enhances nitrogen removal within the B-Stage via carbon-storage driven denitrification even after exposure to aerobic conditions.
- Optimal influent for downstream anammox can be obtained through both NOB out-selection (partial nitritation) and partial denitrification (via internally stored carbon from the sidestream).
- A downstream anammox process allows for low effluent TIN with little-to-no supplemental carbon addition.
- Although sustained partial nitritation and partial denitrification over long-term needs further investigation, the system demonstrates excellent flexibility in being able to operate in both PNA and PdNA modes.
- Future work must bring better understanding of the mechanisms of internal carbon storage for partial denitrification and the impacts of sidestream reactors on AOB/NOB dynamics.

## ACKNOWLEDGEMENTS

HRSD provided the majority of the funding for this work. The authors would like to acknowledge Kathryn Printz, Lindsey Ferguson, Tyler Robinson, and Kaitlyn Greene for their contributions. The authors would also like to acknowledge JangHo Lee, IL Han, Varun Srinivasan, and DongQi Wang who participated in some of the denitrification batch tests and were funded in part by the Water Research Foundation (WRF 4901) and HRSD.

