## Supporting Information for "Sidestream bio-P and mainstream anammox in a BNR process with upstream carbon capture"

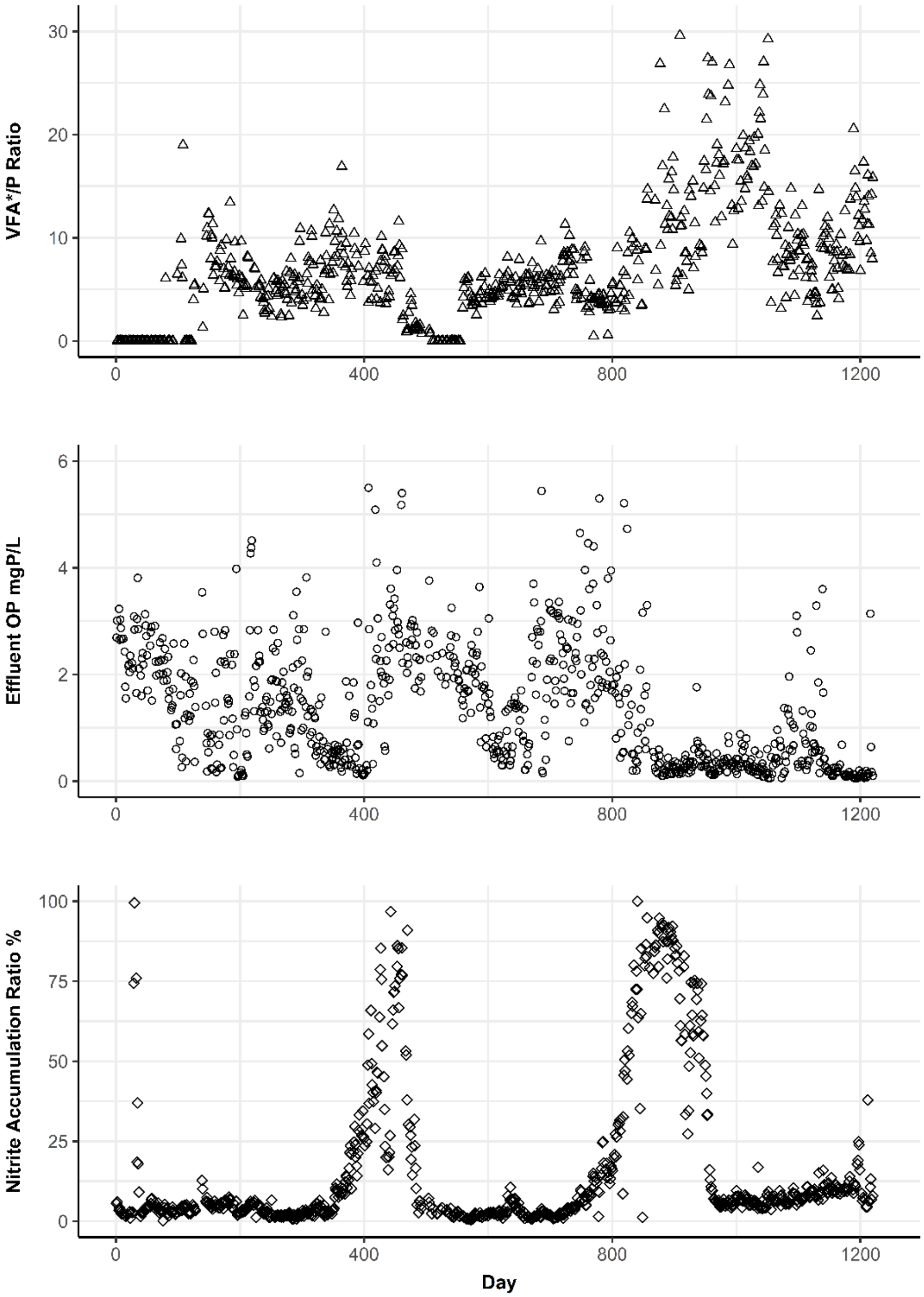


Figure SI-1. Selected operational parameters over the course of the study. Values shown are from 24-hour composite samples.

Table SI-1. Periods of the study and whether successful bio-P and partial nitrification or denitrification occurred in the majority of the phase. Periods provided to generally group times of similar performance or operational conditions in the pilot

| | **Bio-P (effluent P <1 mgP/L)** | **Partial Denitrification** | **Partial Nitrification** |
|---|---|---|---|
| **Period 1** | | | |
| **Period 2** | X | X | |
| **Period 3** | | X | |
| **Period 4** | | | |
| **Period 5** | | | |
| **Period 6** | | | X |
| **Period 7** | X | | X |
| **Period 8** | X | | |

Table SI-1. Selected operational parameters over the course of the study, separated by periods of the study.

| **Period** | **Day Range** | **RAS Split** | **SBPR HRT** | **SBPR SRT** | **Mainstream SRT** | **Sidestream VFA Addition** | **VFA/OP Ratio** | **Fermentate Acetate / Propionate Ratio** |
|---|---|---|---|---|---|---|---|---|
| | | % | hour | day | day | mgVFA*/L | gHAc/gP | gHAc/gPr |
| **1** | 202 – 320 | 21.2 ± 10.1 | 7.6 ± 4.6 | 6.5 ± 5.5 | 8.4 ± 3.0 | 14.4 ± 4.4 | 5.5 ± 1.9 | 1.1 ± 0.4 |
| **2** | 321 – 405 | 28.7 ± 0.7 | 4.1 ± 0.0 | 8.7 ± 3.1 | 8.6 ± 1.9 | 23.8 ± 7.5 | 7.5 ± 2.8 | 1.6 ± 0.7 |
| **3** | 406 – 481 | 28.2 ± 6.0 | 5.3 ± 4.5 | 7.6 ± 3.8 | 7.9 ± 2.0 | 20.3 ± 10.9 | 5.5 ± 2.8 | 1.2 ± 0.5 |
| **4** | 482 – 557 | 7.8 ± 3.3 | 17.8 ± 4.4 | 9.4 ± 3.0 | 12.4 ± 5.1 | 0.6 ± 1.3 | 0.5 ± 0.8 | 1.2 ± 0.4 |
| **5** | 558 – 719 | 32.1 ± 0.6 | 4.1 ± 0.0 | 6.1 ± 1.3 | 9.2 ± 2.5 | 17.9 ± 5.9 | 5.3 ± 1.3 | 1.1 ± 0.3 |
| **6** | 720 – 852 | 31.9 ± 0.0 | 4.1 ± 0.0 | 4.7 ± 1.5 | 5.9 ± 1.6 | 22.1 ± 8.9 | 5.8 ± 2.6 | 1.5 ± 0.5 |
| **7** | 853 – 949 | 31.9 ± 0.0 | 4.1 ± 0.0 | 6.4 ± 2.4 | 8.8 ± 3.7 | 48.9 ± 31.2 | 12.7 ± 5.9 | 0.9 ± 0.2 |
| **8** | 950 - 1220 | 31.9 ± 0.0 | 4.1 ± 0.0 | 5.3 ± 1.9 | 5.4 ± 1.6 | 37.7 ± 16.0 | 12.1 ± 6.1 | 1.1 ± 0.2 |
